# Integration of Diverse Transcriptomics Datasets using Random Forest to Predict Universal Functional Pathways in Tfr Cells

**DOI:** 10.1101/2021.11.29.470410

**Authors:** Alos Diallo, Cecilia B Cavazzoni, Jiaoyuan E Sun, Peter T Sage

**Author notes:** Correspondence (P.T.S.).

## Abstract

**Motivation:** T follicular regulatory (Tfr) cells are a specialized cell subset that controls humoral immunity. Despite a number of individual transcriptomic studies on these cells, core functional pathways have been difficult to uncover due to the substantial transcriptional overlap of these cells with other effector cell types, as well as transcriptional changes occurring due to disease settings. Developing a core transcriptional module for Tfr cells that integrates multiple cell type comparisons as well as diverse disease settings will allow a more accurate prediction of functional pathways. Researchers studying allergic reactions, immune responses to vaccines, autoimmunity and cancer could use this gene set to better understand the roles of Tfr cells in controlling disease progression.

Additional cell types beyond Tfr cells that have similar features of transcriptomic complexity within diverse disease settings may also be studied using similar approaches. High-throughput sequencing technologies allow the generation of large datasets that require specific tools to best interpret the data. The development of a core transcriptional module for Tfr cells will allow investigators to determine if Tfr cells may have functional roles within their biological systems with little knowledge of Tfr biology. With this work, we have addressed the need of core gene modules to define specific subsets of immune cells.

**Results:** We introduce an integrated “core Tfr cell gene module” that can be incorporated into GSEA analysis using various input sizes. The integrated core Tfr gene module was built using transcriptomic studies in Tfr cells from several different tissues, disease settings, and cell type comparisons. Random forest was used to integrate the transcriptomic studies to generate the core gene module. A GSEA gene set was formulated from the integrated core Tfr gene module for incorporation into end-user friendly GSEA. The gene sets are presented along with random genes taken from the GTEX data set and are presented as GMT files. The user can upload the gene set to the GSEA website or any gene set tool which takes GMT files. We also present the full results of the model including p-values calculated by random forest. This provides users with more flexibility in choosing a p-value cutoff that is most appropriate for the experimental setting.

**Availability:** The core Tfr gene sets are freely available at: https://github.com/alosdiallo/TFR_Model. We have also included all of the code and data used in developing these gene sets. The code and results are released under an MIT license.

**Supplementary information:** Supplementary data are available at Bioinformatics online.

## Introduction

Applications of machine learning in immunology and medicine have increased in the past decade(Heng and Painter, 2008) (Eraslan, et al., 2019). These advances have been driven in part by the availability of sizable genomic datasets such as The Genotype-Tissue Expression (GTEX), The Cancer Genome Atlas (TCGA) and Immunological Genome Project (IMMGEN) (Consortium, 2013; Heng and Painter, 2008; McLendon, et al., 2008). The ability to integrate large omics datasets into the training data for machine learning models has also become more prevalent in immunology with the application of tools such as ImmuneML, and CIBERSORT (Newman, et al., 2019; Pavlović, et al., 2021). With these tools and genome wide datasets, researchers have been able to publish custom gene sets for use in gene set enrichment analysis to allow other researchers to leverage this knowledge for their own experiments (Segal, et al., 2004; Subramanian, et al., 2005). Currently, most gene sets and gene modules derived to identify cell types and their functional pathways rely on a single cell type comparison from only one anatomical location and disease state. In this work, we elucidate a core gene module for a specific cell type derived from a wide variety of transcriptomics datasets. We chose the Follicular Regulatory T (Tfr) cell as the subject because of its transcriptional complexity, overlap with related effector cells, presence in multiple tissues, and novel functional roles. This core gene module will serve as a tool for further exploring functional pathways in Tfr cells in immune responses, a cell type that has been associated with controlling disease progression in a number of immunological settings such as vaccination, allergic airway disease, autoimmunity and cancer (Clement, et al., 2019; Eschweiler, et al., 2021; Fu, et al., 2018).

The immune system presents a complex organization, spread throughout the organism, and largely relies on cell-cell communication to achieve protection from infection, to function in tumor suppression, and to maintain a balanced interaction with the microbiota. Antigen recognition is a key feature of the immune system, based on the interaction between B and T cells, which express antigen-specific receptors called BCR and TCR, respectively. The interaction between B and T cells leads to phenotypic changes that correlate with their function. These cells’ activation state, differentiation, and effector function rely on their gene expression changes. Furthermore, factors including genetic factors, drugs and ageing can alter gene expression profiles. These gene expression changes can be assessed by high-throughput sequencing analyses.

Immune responses are finely controlled by the regulatory components of the immune system, such as regulatory T cells (Tregs). They are characterized by the expression of the transcription factor, Foxp3 and can suppress effector T cells through surface molecules and cytokine secretion (Sakaguchi, et al., 2008). T follicular regulatory cells (Tfr) are a transcriptionally and functionally distinct subset of Treg cell. Tfr cells gain access to the B cell follicle within lymphoid organs where they modulate B cell responses and the germinal center reaction (Chung, et al., 2011; Linterman, et al., 2011; Sage and Sharpe, 2020; Wollenberg, et al., 2011). Germinal centers are microanatomical structures where B cells undergo somatic hypermutation and BCR affinity maturation through multiple rounds of interactions with T follicular helper cells (Tfh) (Berek, et al., 1991; Han, et al., 1995; Jacob, et al., 1991; Jacobson, et al., 1974). Tfh cells derive from a pool of mature naïve T cells (conventional T cells) that become activated when their antigen-specific receptor recognizes its antigen along with other stimulatory signals. Tfr cells regulate the germinal center response, enabling the generation of high affinity antibodies while preventing the generation of autoreactive antibodies that could be pathogenic (Clement, et al., 2019; Fu, et al., 2018; Gonzalez-Figueroa, et al., 2021; Lu, et al., 2021; Wu, et al., 2016). Although Tfr cells have been implicated in controlling disease progression in a number of settings, the transcriptional machinery controlling Tfr function remains elusive. This paucity of information is due to the transcriptional complexity of Tfr cells. Tfr cells have a transcriptional program that is a combination of Tfh and Treg cell which is remarkable since neither of these two subsets have the full functionality of Tfr cells (Hou, et al., 2019). Moreover, since Tfr cells can be found in multiple tissues and during multiple disease states, gene expression also correlates with the environment and as well as with functionality. Uncovering a core gene module for Tfr cells that is not dependent on tissue localization or disease setting will allow researchers to narrow focus onto functional pathways that may serve as targets for therapeutics.

Being able to uncover networks of gene expression for individual types of immune cells can help give researchers tools needed to better understand these cells’ phenotype and function. Work from consortiums as well as individual labs has contributed greatly to this goal (Wilk, et al., 2020) (Heng and Painter, 2008). There is also ongoing work to develop an immune cell atlas that may provide a more global view of the immune system. Over the past decade, the development of more sophisticated tools to aid immunologists coupled with decreased sequencing costs have allowed researchers to more effectively develop strategies to examine various cell types in different contexts, such as lymphoid tissues and the tumor microenvironment (Maslova, et al., 2020; Pavlović, et al., 2021). In addition, the use of statistical models in molecular biology can also often provide insight that is difficult or costly to obtain through in vitro experiments (Eraslan, et al., 2019). However, in order to uncover cell type function in large datasets, core modules for cell types need to be generated that are not sensitive to the unique settings of the dataset. To address all of these issues, we developed a core Tfr cell gene module that has been generated using machine learning methods and multiple omics datasets that integrate function, tissue location and disease setting. This core Tfr gene module is designed to be easily used for pathway analysis in the GSEA tool, but can also be used with other gene set enrichment tools such as Panther, and Enrichr (Mootha, et al., 2003) (Subramanian, et al., 2005) (Chen, et al., 2013) (Thomas, et al., 2003). This approach will help the field understand Tfr biology in more detail. In addition, we posit this approach as a key method for generating core transcriptional programs for other cell types that have unique functions but may have complex transcriptional machinery.

## Materials and Methods

There are 77 different polyA-targetted RNA-seq samples taken from mice for this analysis, including Tfh, Tfr, Treg and T conventional cells (naïve T cells) from lymph nodes, spleen, and blood (Figure S1 and supplemental datasets on github). These samples were chosen because they were derived using the same bulk RNAseq library prep pipeline and incorporate effector cells (Tfh, Treg, etc.) with partial overlapping transcriptional signatures as Tfr cells, include multiple tissues and disease states, as well as in vitro studies. The RNA sequencing data in these studies were processed using Qiagen CLC Genomics Workbench version 8 (Qiagen, 2019). Reads were normalized using total exon reads per million (Qiagen, 2019). All of the work on the development of the models and statistics were performed using the R programming language (Team, 2014). Model tuning was done using a training/test set approach with the datasets comprising of bootstrap samples. A training/test gene set was generated from the RNA sequencing data in order to train the statistical models. This data subset is comprised of roughly 2,000 genes based on the following criteria, for Tfr genes: genes had to be expressed two-fold more in Tfr than in TREG for each of the cell types, and genes known to be involved in Tfr biology; for the Non Tfr genes: genes with no expression, genes with two-fold difference in TREG vs TFR for the different cell types, and genes expression below 20% in Tfr cells. We also included genes that have been experimentally validated as having roles in Tfr cells and therefore can serve as a quality control (QC) of the project, for example: CXCR5, ICOS, PD-1, CCR7, Sell, and the recently described Nrn1 (Chung, et al., 2011; Gonzalez-Figueroa, et al., 2021; Linterman, et al., 2011; Wollenberg, et al., 2011). In addition, we assessed correlation between genes from the two classes to determine whether the statistical models would likely model noise as opposed to a core Tfr transcriptional module (Figure S2). To understand whether the two classes were highly correlated, as well as how correlated the genes within the classes were, Pearson correlation was performed on the training and test data (Figure S3).

In order to determine which statistical method would best model the core Tfr gene module we generated 200 bootstrap samples and gauged based on metrics from the ROCR package (Sing, et al., 2005). For a given bootstrap sample, the data was divided into a training and test set with a 70:30 split, respectively, using the sample function. Each sample was randomly generated, which allows for the training and test data to be independent of one another. Each individual training and test sample was then used to fit random forest (RF), linear discriminant analysis (LDA), naïve bayes (NB), K nearest neighbors (KNN), Support Vector Machine (SVM), and logistic regression (LR) (Wiener, 2002) (Jerome H. Friedman, 2010) (David Meyer, 2021) (Venables, et al., 2002). This was carried out to test a range of statistical models to approach the problem using several different strategies. We looked at several different approaches to be able to cover a wide range of methods, including Bayesian, algorithmic, and clustering based approaches to determine which strategy would be most effective at classifying our samples. Results were compared using error rate, accuracy, specificity, and sensitivity, all of which were calculated in the ROCR package (refer to Supplementary methods) (Sing, et al., 2005).

Once all of the different statistical models were tested, RF was chosen as the model for further analysis based on superior results during performance tests (Figure 1). Once we determined that random forest should be used based on the performance metrics from figure 1, we then assessed if the model was prone to bias that could skew the results. To do this, we examined the effects of different variables in the training versus test set (Figure S4). In this analysis, three different datasets were used to fit random forest. First, the training and test sets contained the same column information (variables) with only the genes being randomly selected (Figure S4). Second, the columns used were randomly selected for the training set and test set with the columns used for the selection of each being independent from one another (Figure S4). We then examined the effects of changing the class designation of genes in the training and test sets (Figure S5). In this case, genes were randomly assigned to one of the two classes and used to fit the random forest model. This was compared with a dataset where the class designations were not altered (Figure S5).

**Figure 1:**
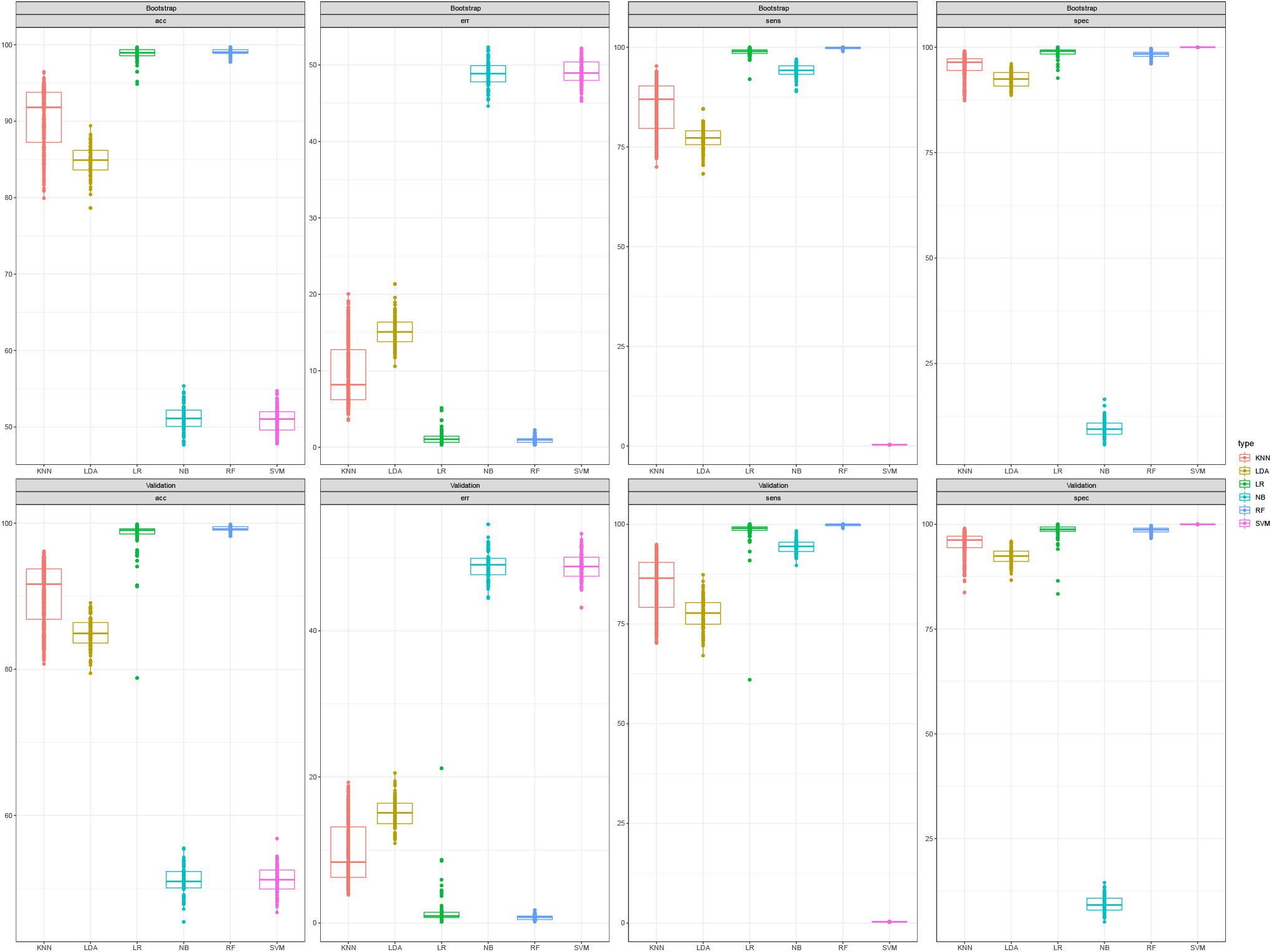
Performance Metrics for each of the Statistical Models used Boxplots depicting performance metrics on accuracy, error, sensitivity, and specificity were calculated for KNN, LDA, LR, NB, RF, and SVM. Each boxplot represents 100 bootstrap samples.

In order to optimize RF, we assessed different model parameters. The parameter mtry in the random forest model were tuned using the tuneRF function, which outputs the proportion of variables used when building trees in RF, and ntreeTry which indicates the number of trees to generate (Wiener, 2002). This was tested on 300 trees (300 was chosen because the parameters are stabilized at this number). In other words, the parameter mtry remained constant when the number of trees exceeded 300 trees. While using tuneRF, values from 1 to 36 were chosen for the parameter of mtry, as values above 36 did not yield an increase in performance. An mtry value of 18 was chosen in order to obtain the lowest training error, with the lowest model complexity. The final random forest model was then run against the whole dataset to generate the core Tfr gene module. All of the plots in this analysis are created using these functions: ggplot, corplot, partialPlot, tuneRF (Wickham, 2016) (Wiener, 2002) (Revelle, 2021).

The top 2,000 ranking genes in the core Tfr gene module produced by RF were made into a custom gene set to be used in common bioinformatics pipelines (See top 2000 gene set in Supplementary data). These gene lists are stored as rows of 200 genes in a single “.gmt” file for use with the Broad Institute’s GSEA tool (Reich, et al., 2006). These can be used in other bioinformatics tools which use a “.gmt” file as input. We also included 2,000 random genes used as a null gene set model, this was used as a negative control to assess scores of random genes. For this, this we picked the random (using the sample function in R) genes from the GTEX dataset that comprise 20 individual gene sets (Consortium, 2013). GTEX was chosen as the source for the NULL model gene sets for 3 reasons. First, the GTEX analysis was conducted using healthy tissue (non-tumor). Second, the data came from many individual patients which allows for great variability in samples (1,000 individuals). Finally, the GTEX data is made up of samples from 54 different tissue types (Consortium, 2013). This strategy ensures the NULL model is not heavily impacted by Tfr gene expression, as many tissues will likely not contain large populations of immune cells. Using the GSEA tool, the custom gene sets (for GSEA) were then run against the subsets of the original RNASeq data to determine the enrichment for Tfr gene expression. We compared samples from Tfr, vs Treg cells as well as Tfr vs Tcon, and Tfr vs Tfh cell samples from each of the different tissues. The final QC analysis performed using the GTEX data was to use the genes from the training and test set, but with data from the different GTEX tissues (Figure S6). The data was randomly sampled (Figure S6). This was to determine if RF would model noise as well as it could model the normal training and test data (Figure S6).

All of the RNA sequencing data, along with the custom gene sets are stored on github, in addition, the code used in this project is also stored on github at https://github.com/alosdiallo/TFR_Model.

## Results

The results of comparing the training and test errors for the bootstrap samples shown in the boxplots of each of the models shows that random forest had the lowest median error, whereas naïve bayes had the highest error (see Figure 1 and Table 1), according to all 4-performance metrics: accuracy, error rate, sensitivity, and specificity (See Supplemental text for math). Table 1 shows that random forest had the lowest error rate, highest accuracy, highest sensitivity, and second highest specificity. The second-best performing method was LR followed by KNN, LDA, SVM, and finally NB (see Figure 1, and Table 1). Figure 1 shows the results of all 200 bootstrap samples. This was broken up into 2 separate batches, where 100 bootstrap samples were built and tested for each batch (Figure 1). LDA had the largest distribution in error for the 200 samples, with RF having the tightest spread, which points to RF being the most reproducible of the methods tested (Figure 1). Based on these results, we further focused on random forest as a strategy since it had the lowest error rate and highest accuracy which more appropriately fit within the goals of the project.

**Table 1:** Performance of Statistical Models

| Method | Accuracy | Error Rate | Sensitivity | Specificity |
| --- | --- | --- | --- | --- |
| Logistic regression (LR) | 0.98156231 | 0.01831329 | 0.97874395 | 0.9863281 |
| Random forest (RF) | 0.99034749 | 0.01074219 | 0.9979188 | 0.9838188 |
| Naïve Bayes (NB) | 0.5135401 | 0.48703125 | 0.94472021 | 0.090625 |
| Linear discriminant analysis (LDA) | 0.838853 | 0.1640048 | 0.7617178 | 0.9168317 |
| K Nearest Neighbors (KNN) | 0.9141252 | 0.08427 | 0.8655876 | 0.9635762 |
| Support Vector Machine (SVM) | 0.5104334 | 0.4895666 | 0.003267974 | 1 |

Median Performance metrics: accuracy, error, sensitivity and specificity for the different statistical models. The models used for this analysis are LR, RF, NB, LDA, and KNN.

After running the model against the full dataset, roughly 5,000 genes were predicted by the model to be associated with the core Tfr gene module (Figure 2A). The top 2,000 genes were chosen to be used in the gene module based on their p-value (0.77 for Tfr). A p-value of 0.77 was chosen because it provided a good balance between a conservative model while also including a large sample of genes. We have included the full list (of 5,000 genes) along with their p-values to allow users to alter the cutoff should they choose to be more or less conservative. Examining the variable importance plot, we can see that Lymph node Tfr (LNTfr) samples (see Figure 2B) have the largest effect on the model performance followed by the lymph node Treg samples (see Figure 2B). Lymph node was followed by spleen, then blood in terms of importance as predicted by the model (Figure 2B). Overall, the goal was to be able to integrate diverse transcriptomics datasets to be able to predict functional pathways which can represent a proxy for Tfr gene expression, but we therefore also have a model for functional pathways that exclude Tfr gene expression. Samples with Tfr cells had the highest association with the Tfr model whereas Treg samples were most highly associated with not being affiliated with Tfr gene expression (Figure 2B). This confirms the ability of the model to predict differences, since Tfr cells derive from natural Tregs but are phenotypically and functionally distinct (Sage and Sharpe, 2020). Additionally, we compared genes of known association with specific T cell phenotypes which were predicted to be related to Tfr gene expression by the model, such as *Cxcr5, Icos* and *Pdcd1*. (Figure 3A). The results show that the most relevant genes for the Tfr model include a selection of genes that are known to be characteristic of this T cell subset, and when the expression of such genes is compared to Treg and Tfh cell samples, Tfr samples cluster together as opposed to Treg and Tfh samples (Figure 3A). This was the case for lymph node, spleen, and blood samples (Figure 3A). These results serve to further validate the model.

**Figure 2:**
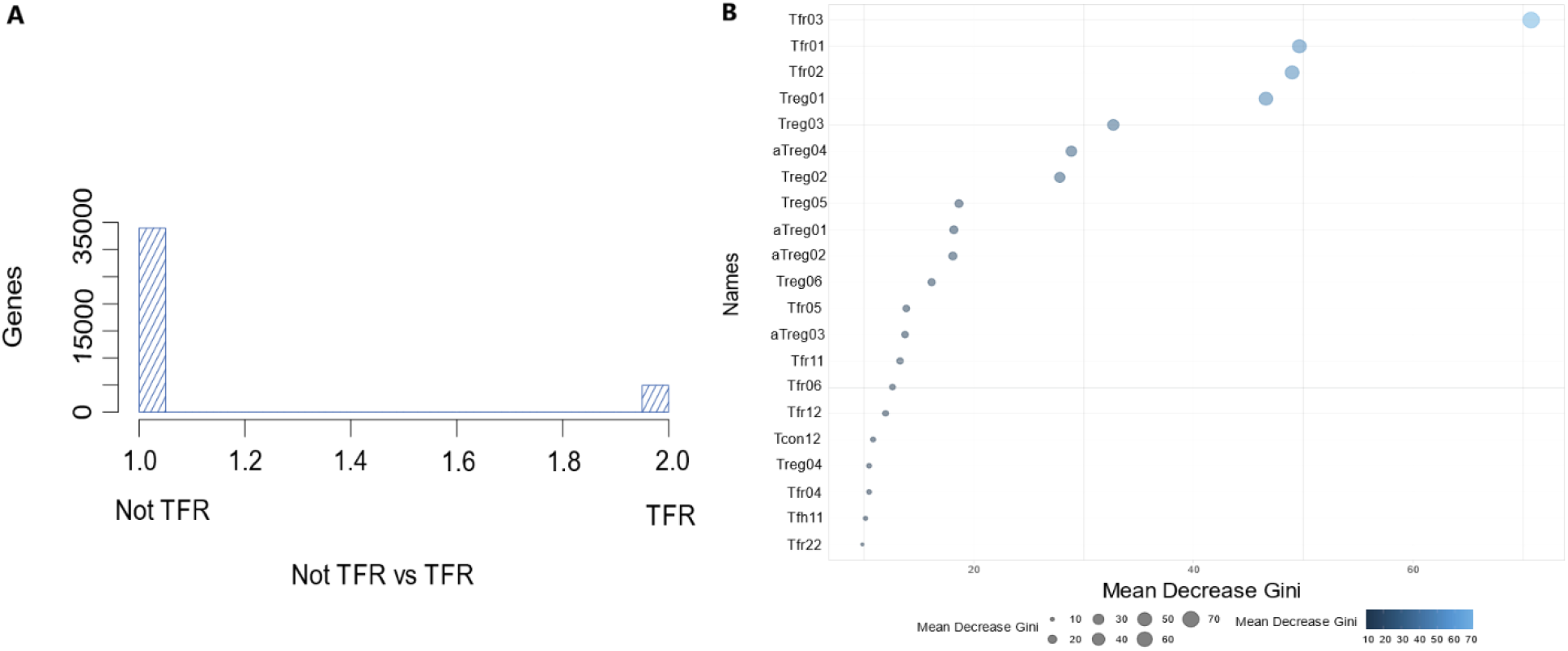
Model Selection and Importance A: Results of the model selection by random forest based on the two classes, Tfr and the null model. B: Variable importance of datasets from applying random forest to the test set.

**Figure 3:**
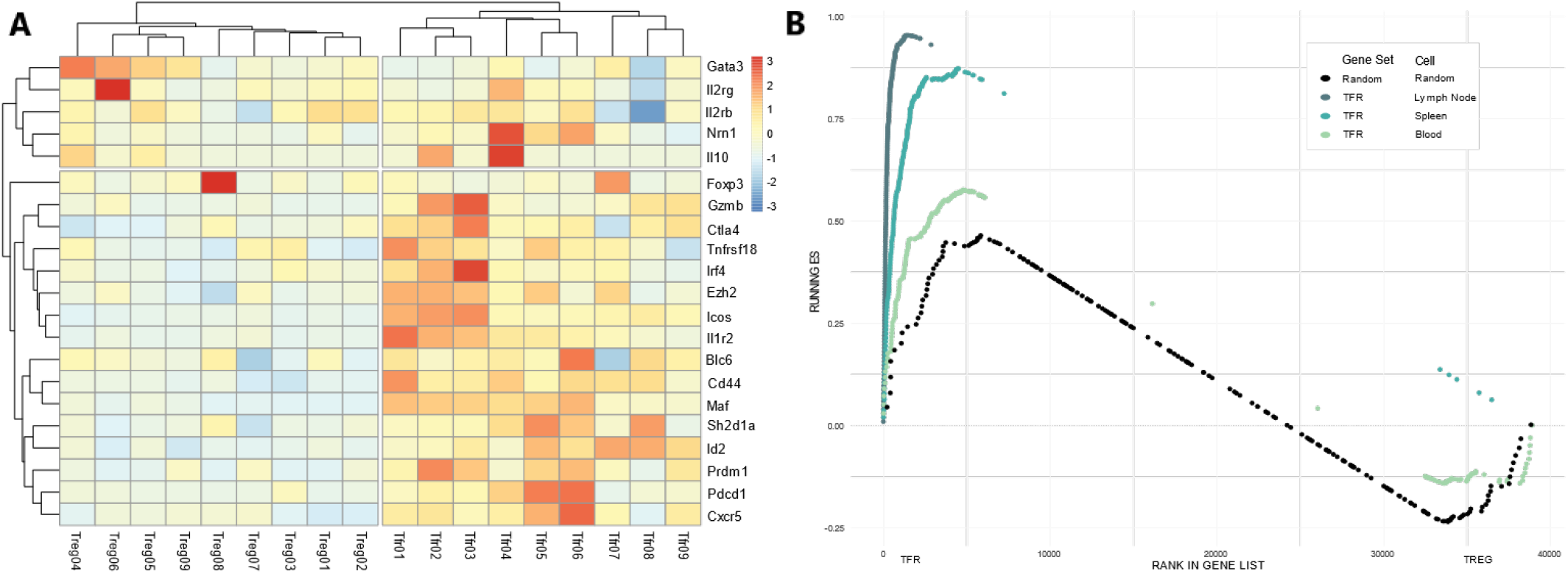
Model results A: Heatmap showing genes known to be associated with the Tfr cell phenotype that were selected by the model as being TFR related. B: GSEA results for the custom gene set and the random gene sets run against the TFR and TREG datasets.

Corroborating the previous result, we observed that the core Tfr gene module when applied to Tfr and Treg samples from lymph nodes, spleen and blood were able to enrich using gene set enrichment analysis (GSEA) for Tfr cells in the Tfr samples as compared to Treg samples. On the contrary, random gene sets generated from the GTEX database applied to Tfr and Treg samples from lymph nodes, spleen and blood showed no particular enrichment or favorability toward either cell type (see Figure 3B). This is significant as it supports our hypothesis that the predicted gene list is specifically involved in Tfr cell gene expression. These results are observed in the lymph node, spleen and blood samples (Figure 3B). Blood, in this case shows the least enrichment of the three tissue types. This is aligned with our expectations since Tfr cells in the blood present distinct functionality and memory-like features that are not shared with Tfr cells in lymphoid organs (Sage, et al., 2014) (Figure 3B).

## Discussion and Conclusion

A main goal of this work is to provide a gene module that can be used by immunologists to better identify Tfr cells in big datasets and to provide a tool to understand the role of Tfr cells in their experiments. The other goal is to develop a procedure that can be implemented in order to characterize other immune cells. Similar work has been done for PD-1 expression (Diallo, 2021). We were able to identify roughly 2,000 genes which were found to contribute to the Tfr gene expression profile and show how statistical models like random forest can be used in modeling immune cells (Figure 2A). Within the top 2,000 genes, we were able to identify several previously described genes that are specifically related to Tfr function, such as *Cxcr5, Icos* and *Pdcd1* as well as the recently described *Nrn1* (Figure 3A). The results of running random forest against the bootstrap samples showed very consistent results (Figure 1). This likely means that random forest could work well for different immune cell types. Using tools and larger databases such as IMMGEN and ImmuneML, it should be possible to build up a database for modeling more cell types (Heng and Painter, 2008; Pavlović, et al., 2021).

In terms of the other models that were tested, KNN shows potential, but likely researchers would have to experiment with different distance measures. This has been shown to work well in other instances (McDermott, et al., 2020; Parry, et al., 2010). The results found with random forest were consistent with what we previously found for Tfr cell gene expression when compared with Tfh and Treg cells (Hou, et al., 2019). This is particularly relevant since Tfr cells share features with both Tfh and Treg cells and were still properly separated from these T cell types. For instance, in figure 2B, the lymph node Tfr and lymph node Treg samples played the largest role in characterizing the model, which is in agreement with the biology of these cell types. Secondary lymphoid organs, such as lymph nodes, are an important site of activity of these T cells subsets where their effector and suppressive functions fully develop. Looking at the variable importance plot for RF, Tfr cells were the most important for the Tfr model and Treg samples were the most important for the null model. This is what we expected to find, as we have previously observed that the transcriptome of Tfr cells are more distant from Treg cells than from Tfh cells (Hou, et al., 2019). The heatmap in figure 3A also shows good cluster separation between the different cell types for the models associated with Tfr cell gene expression. In all the analyses performed with the Tfr model, it is possible to observe the distinction between Tfh, Treg and Tfr cells which is very important due to the sharing of some features of Tfr cells with Treg and Tfh cells that could represent confusion factors in gene expression analyses. Finally, we also noticed that when testing the gene set against Tfr and Treg samples, enrichment was heavily skewed towards Tfr cell gene expression. This was in stark contrast to the results seen using the GTEX sample data. We would expect this if our gene set was able to positively characterize Tfr cell gene expression in part because it differs so much from the negative control. Therefore, these results confirm the ability of the model to identify Tfr cell gene expression which does not happen when applied to random gene sets.

Tfr cells have been shown to play important roles in modulating germinal center reactions, controlling antibody responses to vaccination and allergic immune responses (Clement, et al., 2019; Fu, et al., 2018; Gonzalez-Figueroa, et al., 2021; Lu, et al., 2021; Wu, et al., 2016). Moreover, in aged mice, increased frequency of Tfr cells contributes to impaired antibody response after immunization (Sage, et al., 2015). These cells have also been recently implicated in impaired antitumor immunotherapy (Eschweiler, et al., 2021). In humans, altered Tfr cell frequency in peripheral blood has been associated with more severe COVID-19 disease, as well as with autoimmune diseases, such as lupus (Gong, et al., 2020; Xu, et al., 2017). Additionally, increased frequencies of Tfr cells have been found in tumor-draining lymph nodes and peripheral blood from patients with different types of cancer (reviewed in (Huang, et al., 2020). Tfr cells have also been implicated in the persistence of HIV (Miller, et al., 2017). Therefore, a better understanding of Tfr cell biology and the immunoregulation promoted by these cells is of interest and may have implications in many clinical settings.

The use of computationally derived gene sets can allow for the ability to leverage much of the genomic data that exists in the public domain. In the future, it should be possible to leverage other types of genomic data such as single cell, or HiC experimental data to further enhance the usefulness of such machine learning models. We also provide a framework for carrying out this sort of work for other cell types using bulk RNA Sequencing data. Other tools such as language models may also play a role in more accurately modeling biology. In summary, we have presented a Tfr cell gene expression gene list that the authors hope will help further research by providing new insights into how Tfr cells could impact their work.

## Data availability

Data is available from the senior author upon reasonable request.

## Acknowledgments

We would like to thank Victor Farutin for his feedback on the model’s development.

