## Supplementary material for "Integration of Diverse Transcriptomics Datasets using Random Forest to Predict Universal Functional Pathways in Tfr Cells": S1,S2,S3, S4, S5, S6, Supplemental Method

Supplementary Information
### The Development of a TFR Gene Expression Data Set using Random Forest
Alos Diallo, Cecilia B Cavazzoni, Jiaoyuan E Sun, Peter T Sage

| Name | Cell Type | Tissue | Disease State | Vivo/Vitro | Accession |
|---|---|---|---|---|---|
| aTreg | ICOS+ Treg | LN | Vaccination | Ex Vivo | GSE124884 |
| aTreg | ICOS+ Treg | LN | Vaccination | Ex Vivo | GSE124884 |
| aTreg | ICOS+ Treg | LN | Vaccination | Ex Vivo | GSE124884 |
| aTreg | ICOS+ Treg | LN | Vaccination | Ex Vivo | GSE124884 |
| ExTfr01 | ExTfr | LN | Vaccination | In Vitro | GSE124884 |
| ExTfr02 | ExTfr | LN | Vaccination | In Vitro | GSE124884 |
| ExTfr03 | ExTfr | LN | Vaccination | In Vitro | GSE124884 |
| Tcon01 | Tconv | LN | Vaccination | Ex Vivo | GSE124884 |
| Tcon02 | Tconv | LN | Vaccination | Ex Vivo | GSE124884 |
| Tcon03 | Tconv | LN | Vaccination | Ex Vivo | GSE124884 |
| Tcon04 | Tconv | Spleen | Vaccination | Ex Vivo | GSE124884 |
| Tcon05 | Tconv | Spleen | Vaccination | Ex Vivo | GSE124884 |
| Tcon06 | Tconv | Spleen | Vaccination | Ex Vivo | GSE124884 |
| Tcon07 | Tconv | Blood | Vaccination | Ex Vivo | GSE124884 |
| Tcon08 | Tconv | Blood | Vaccination | Ex Vivo | GSE124884 |
| Tcon09 | Tconv | Blood | Vaccination | Ex Vivo | GSE124884 |
| Tcon10 | Tconv | LN | Vaccination | Ex Vivo | GSE134153 |
| Tcon11 | Tconv | LN | Vaccination | Ex Vivo | GSE134153 |
| Tcon12 | Tconv | LN | Vaccination | Ex Vivo | GSE124884 |
| Tcon13 | Tconv | LN | Vaccination | Ex Vivo | GSE124884 |
| Tcon14 | Tconv | LN | Vaccination | Ex Vivo | GSE124884 |
| Tcon15 | Tconv | LN | Vaccination | Ex Vivo | GSE124884 |
| Tfh01 | Tfh | LN | Vaccination | Ex Vivo | GSE124884 |
| Tfh02 | Tfh | LN | Vaccination | Ex Vivo | GSE124884 |
| Tfh03 | Tfh | LN | Vaccination | Ex Vivo | GSE124884 |
| Tfh04 | Tfh | Spleen | Vaccination | Ex Vivo | GSE124884 |
| Tfh05 | Tfh | Spleen | Vaccination | Ex Vivo | GSE124884 |
| Tfh06 | Tfh | Spleen | Vaccination | Ex Vivo | GSE124884 |
| Tfh07 | Tfh | Blood | Vaccination | Ex Vivo | GSE124884 |
| Tfh08 | Tfh | Blood | Vaccination | Ex Vivo | GSE124884 |
| Tfh09 | Tfh | Blood | Vaccination | Ex Vivo | GSE124884 |
| Tfh10 | Tfh | LN | Vaccination | In Vitro | GSE124884 |
| Tfh11 | Tfh | LN | Vaccination | In Vitro | GSE124884 |
| Tfh12 | Tfh | LN | Vaccination | In Vitro | GSE124884 |
| Tfh13 | Tfh | LN | Vaccination | Ex Vivo | GSE134153 |
| Tfh14 | Tfh | LN | Vaccination | Ex Vivo | GSE134153 |
| Tfh15 | Tfh | LN | Allergy | Ex Vivo | GSE134153 |
| Tfh16 | Tfh | LN | Allergy | Ex Vivo | GSE134153 |
| Tfr01 | Tfr | LN | Vaccination | Ex Vivo | GSE124884 |
| Tfr02 | Tfr | LN | Vaccination | Ex Vivo | GSE124884 |
| Tfr03 | Tfr | LN | Vaccination | Ex Vivo | GSE124884 |
| Tfr04 | Tfr | Spleen | Vaccination | Ex Vivo | GSE124884 |
| Tfr05 | Tfr | Spleen | Vaccination | Ex Vivo | GSE124884 |
| Tfr06 | Tfr | Spleen | Vaccination | Ex Vivo | GSE124884 |
| Tfr07 | Tfr | Blood | Vaccination | Ex Vivo | GSE124884 |
| Tfr08 | Tfr | Blood | Vaccination | Ex Vivo | GSE124884 |
| Tfr09 | Tfr | Blood | Vaccination | Ex Vivo | GSE124884 |
| Tfr10 | Tfr | LN | Vaccination | Ex Vivo | GSE124884 |
| Tfr11 | Tfr | LN | Vaccination | Ex Vivo | GSE124884 |
| Tfr12 | Tfr | LN | Vaccination | Ex Vivo | GSE124884 |
| Tfr13 | Tfr | LN | Vaccination | In Vitro | GSE124884 |
| Tfr14 | Tfr | LN | Vaccination | In Vitro | GSE124884 |
| Tfr15 | Tfr | LN | Vaccination | In Vitro | GSE124884 |
| Tfr16 | Tfr | LN | Vaccination | Ex Vivo | GSE134153 |
| Tfr17 | Tfr | LN | Vaccination | Ex Vivo | GSE134153 |
| Tfr18 | Tfr | LN | Allergy | Ex Vivo | GSE134153 |
| Tfr19 | Tfr | LN | Allergy | Ex Vivo | GSE134153 |
| Tfr20 | Tfr | LN | Vaccination | Ex Vivo | GSE124884 |
| Tfr21 | Tfr | LN | Vaccination | Ex Vivo | GSE124884 |
| Tfr22 | Tfr | LN | Vaccination | Ex Vivo | GSE124884 |
| Tfr23 | Tfr | LN | Vaccination | Ex Vivo | GSE124884 |
| Treg01 | Treg | LN | Vaccination | Ex Vivo | GSE124884 |
| Treg02 | Treg | LN | Vaccination | Ex Vivo | GSE124884 |
| Treg03 | Treg | LN | Vaccination | Ex Vivo | GSE124884 |
| Treg04 | Treg | Spleen | Vaccination | Ex Vivo | GSE124884 |
| Treg05 | Treg | Spleen | Vaccination | Ex Vivo | GSE124884 |
| Treg06 | Treg | Spleen | Vaccination | Ex Vivo | GSE124884 |
| Treg07 | Treg | Blood | Vaccination | Ex Vivo | GSE124884 |
| Treg08 | Treg | Blood | Vaccination | Ex Vivo | GSE124884 |
| Treg09 | Treg | Blood | Vaccination | Ex Vivo | GSE124884 |
| Treg10 | Treg | LN | Vaccination | Ex Vivo | GSE124884 |
| Treg11 | Treg | LN | Vaccination | Ex Vivo | GSE124884 |
| Treg12 | Treg | LN | Vaccination | Ex Vivo | GSE124884 |
| Treg13 | Treg | LN | Vaccination | Ex Vivo | GSE124884 |
| Treg14 | Treg | LN | Vaccination | Ex Vivo | GSE124884 |
| Treg15 | Treg | LN | Vaccination | Ex Vivo | GSE124884 |
| Treg16 | Treg | LN | Vaccination | Ex Vivo | GSE124884 |

Supplemental Figure S1: List of RNAseq datasets used in model generation.

Supplemental Methods:

In order to compare statistical models, specific performance metrics were chosen. Results are compared using error rate $P(\hat{Y} \neq Y)$, which is defined as: $\frac{FP+FN}{P+N}$, accuracy: $P(\hat{Y} = Y)$, $\frac{TP+TN}{P+N}$, specificity: $P(\hat{Y} = \ominus \mid Y = \ominus)$, and sensitivity: $P(\hat{Y} = \oplus \mid Y = \oplus)\ \frac{TP}{P}$, all of which came from the ROCR package {Sing, 2005 #904}.

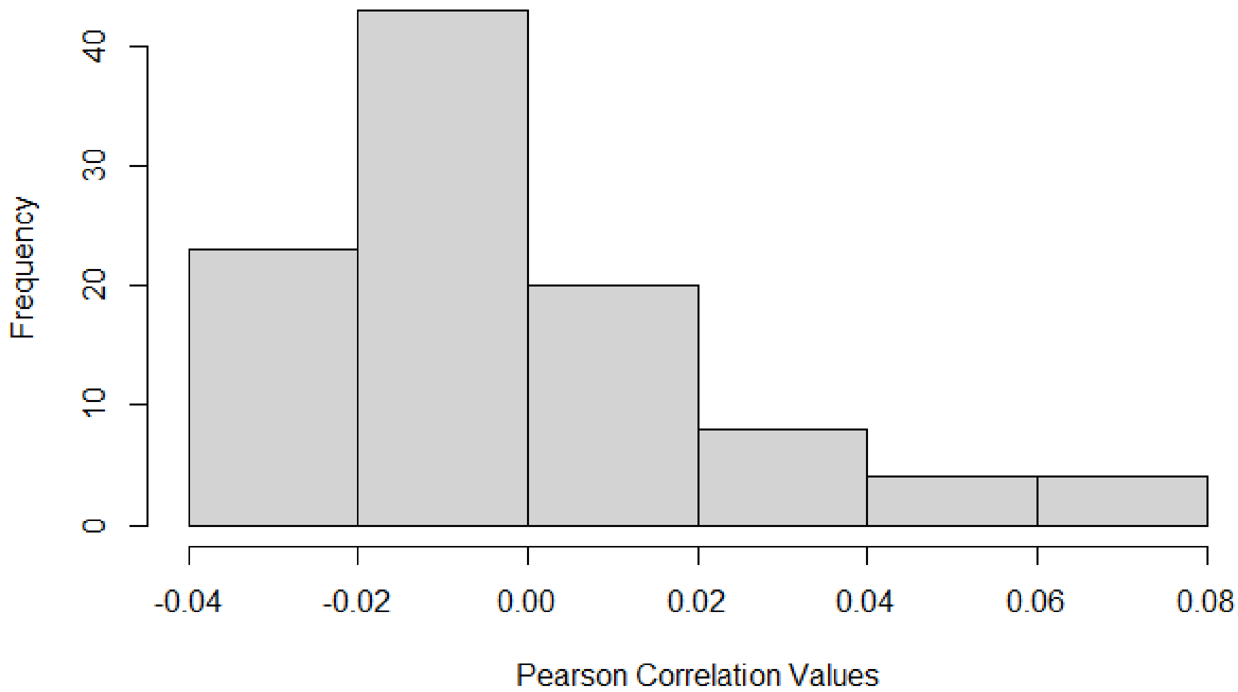


Supplemental Figure S2: Histogram of TFR vs Non-TFR Data. A histogram showing how correlated data are for the two class categories in the training/test set. This indicates that the data from the two classes are not highly correlated.

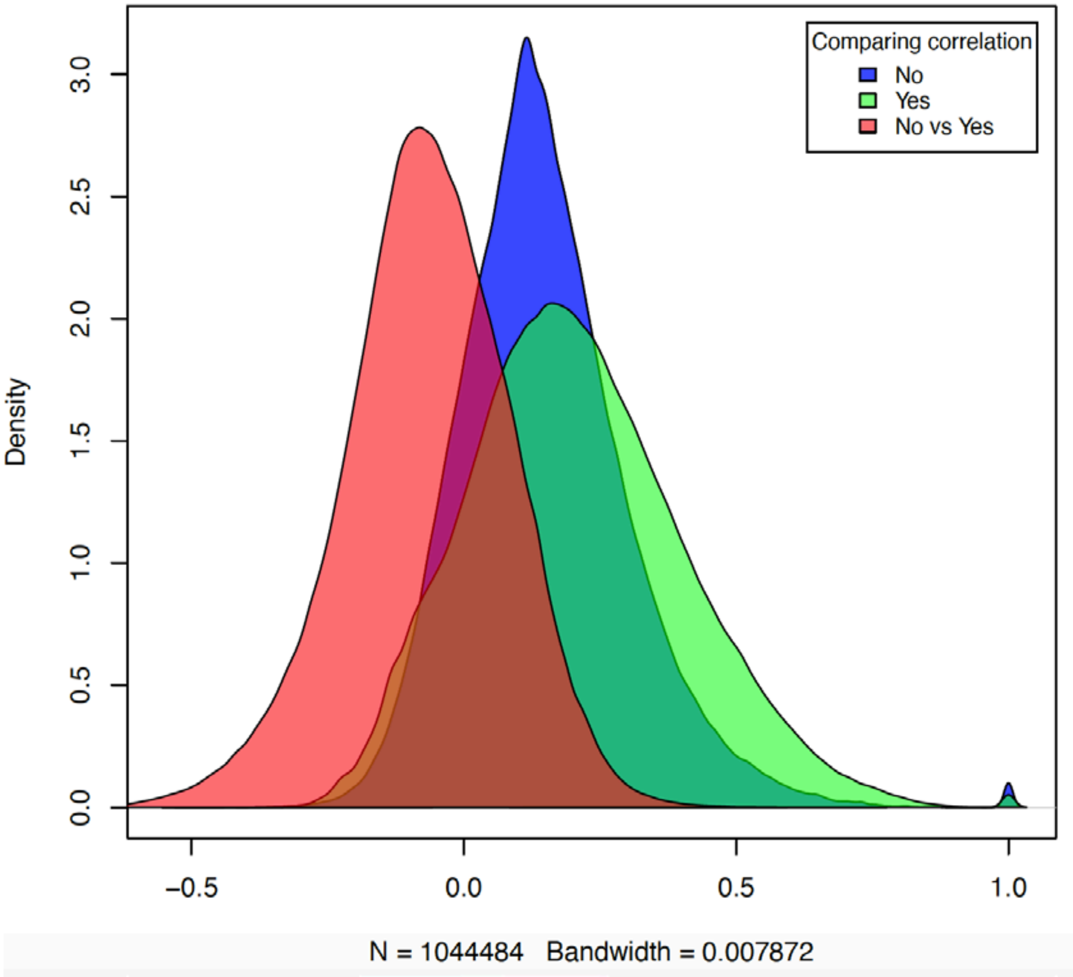


Supplementary Figure S3: Correlation of gene expression data for a given class. No are genes not associated with the TFR model, Yes are genes which are associated with the TFR model, and “No vs Yes” contains all genes. These results indicate that the model is not simply modeling the correlations in the data.

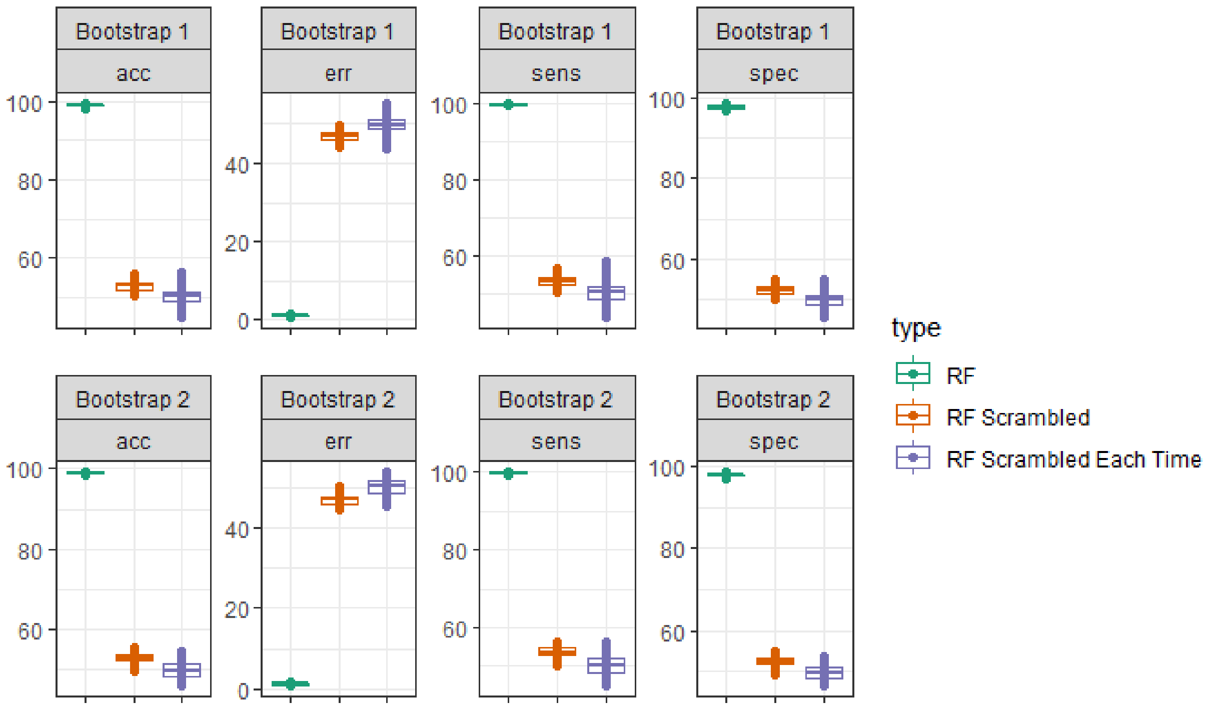


Supplemental Figure S4: A test to determine if the column assignments effect model performance. Comparing 200 bootstrap samples for Random Forest under different conditions RF: where all of the variables (columns) are fed into the Random Forest algorithm, RF Scrambled: where different columns were chosen for the training vs the test set, RF Scrambled Each Time: where different columns were chosen for each bootstrap sample. In the case of “RF Scrambled” and “RF Scrambled each time” the columns used for the training and test set are chosen at random and therefor variables used in the training set are not guaranteed to be in the test set. These results indicate that the variables (columns) used to training the data must also be used to test the data.

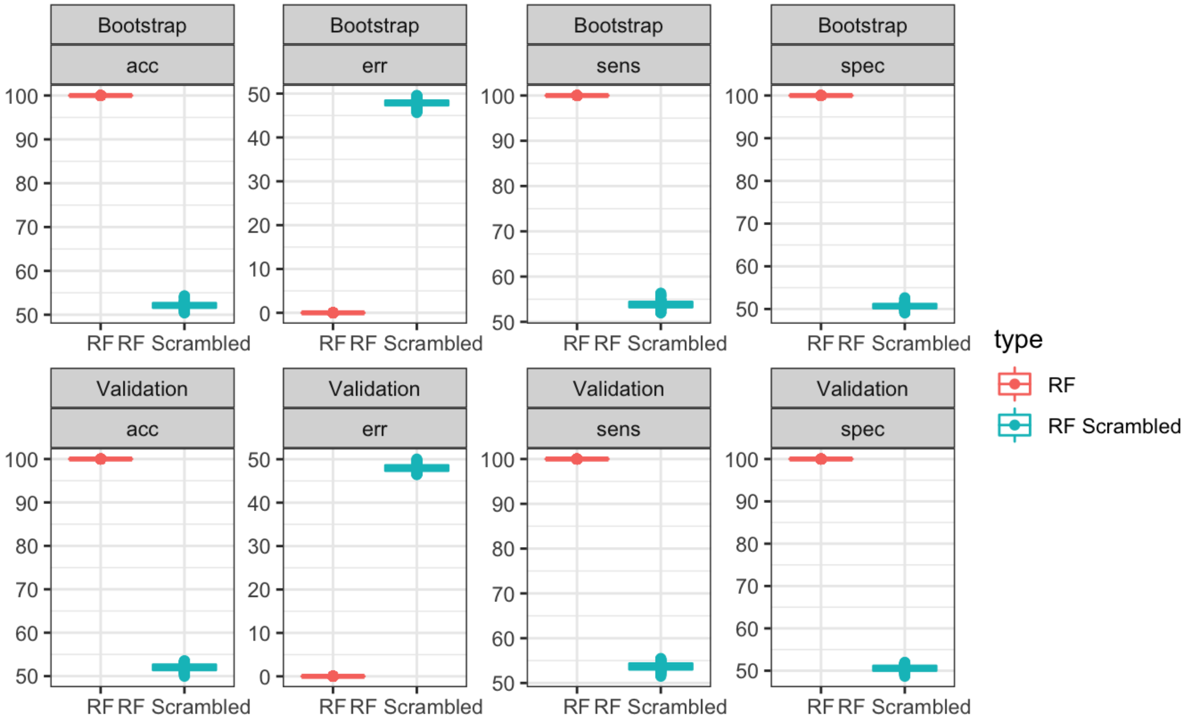


Supplementary Figure S5: Randomly scrambling the class designations. Results from 200 bootstrap samples (models) in which we run random forest on the training and test set. The plot shows data where the class designation is normal (RF), versus data where the class designations were randomly scrambled (RF Scrambled). This means that the rows used for the TFR class vs NULL were changed randomly. The results of this indicates that the genes used to model TFR expression are important. This is because the model is not able to make accurate predictions when the row assignments are scrambled.

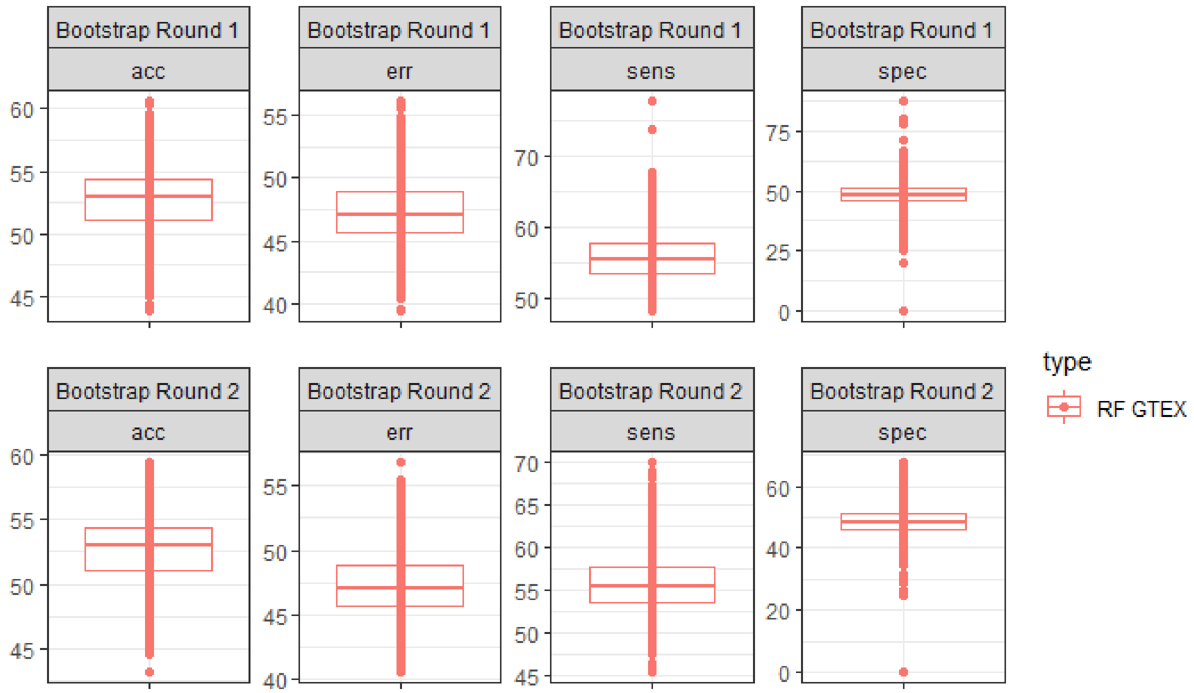


Supplemental Figure S6: Results of random forest using gene expression values from GTEX. The results of 200 bootstrap samples where the data comes from GTEX data, and the gene names are from the training/test set. This indicates that the genes used in the training and test set are important to the model’s ability to make accurate predictions.
